## Supplementary Fig. 1; Supplementary Fig. 2; Supplementary Fig. 3; Supplementary Fig. 4; table S1 for "Exploration of Mechanisms of Drug Resistance in a Microfluidic Device and Patient Tissues"

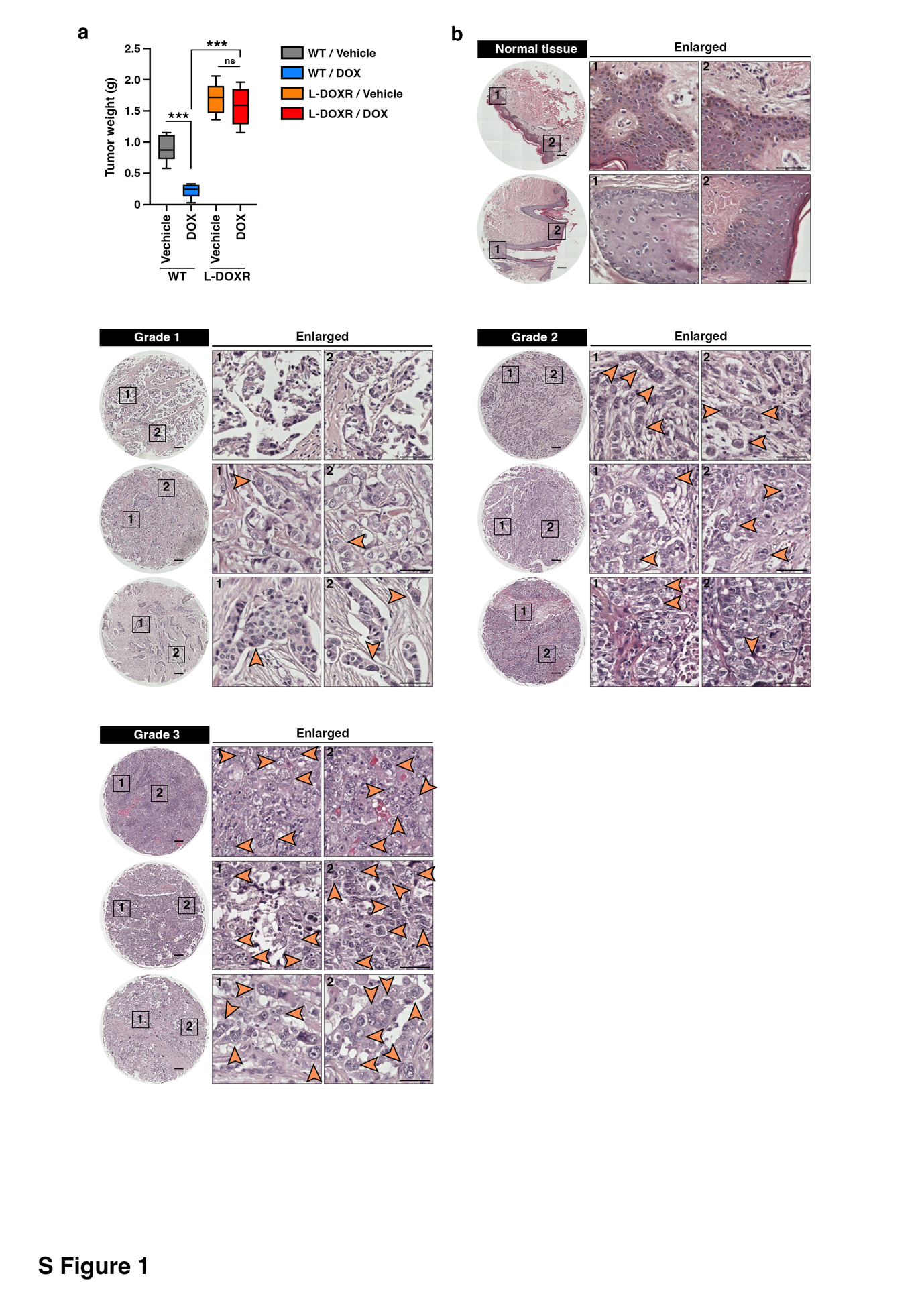

**Supplementary Fig. 1 (related to Fig. 3). L-DOXR accelerated cancerous growth and tumor progression in TNBC.** (a) Wet weights of WT and L-DOXR tumors treated with DOX or vehicle. (b) Representative images showed triple-negative breast cancer tissue microarray (TMA, US Biomax #BR1301) with different tumor grades (grade 1, 2, 3, and negative) after stained with H&E (Hematoxylin and eosin) for the detection of L-DOXR. Black boxes are magnified. Orange arrows indicate PACCs. Scale bar: 500 μm.

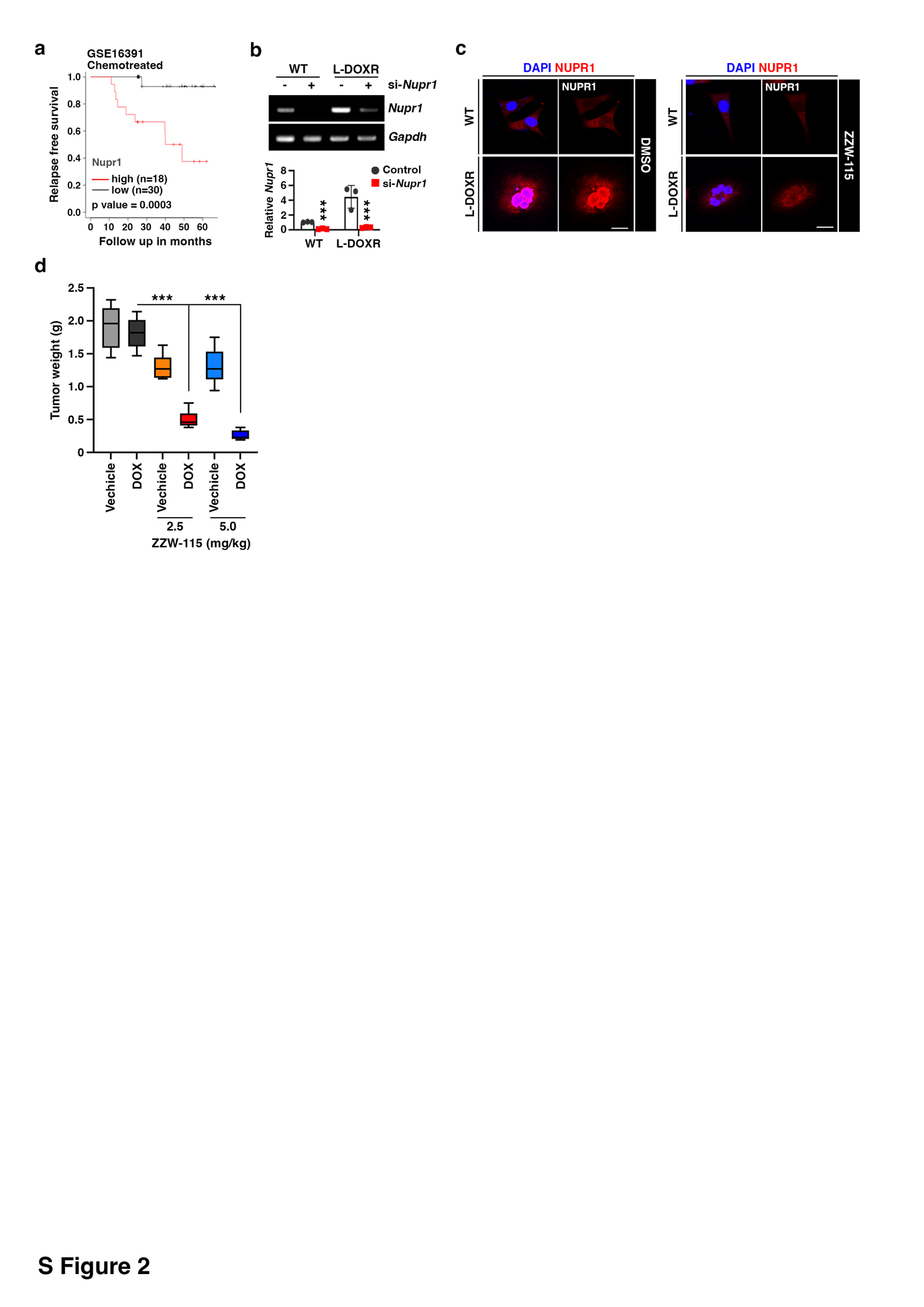

**Supplementary Fig. 2 (related to Fig. 4). NUPR1 is a key mediator of chemoresistance in L-DOXR.** (a) Comparison of overall survival between high (red line) or low (blue line) expressions of *Nupr1* in patients after chemo treated (GSE16391) were analyzed by the Kaplan-Meier and Log-rank test. KM survival curve representing the relapse-free survival rate in chemotherapy-treated patients (n=48) based on low (n=30) vs. high (n=18) *Nupr1* expression. (b) si-*Nupr1* was transfected into WT and L-DOXR and incubated for 24h. The RNA expressions of *Nupr1* were measured by RT-PCR. (c) WT and L-DOXR were stained with anti-NUPR1 (red) after treated ZZW-115. (d) Tumor weights were measured after they were harvested from the mice. Scale bar: 20 μm. All data are presented as means ± SEM; ***p < 0.001; Student’s two-tailed, unpaired t-test (b); one-way ANOVA with Bonferroni’s post-test (d).

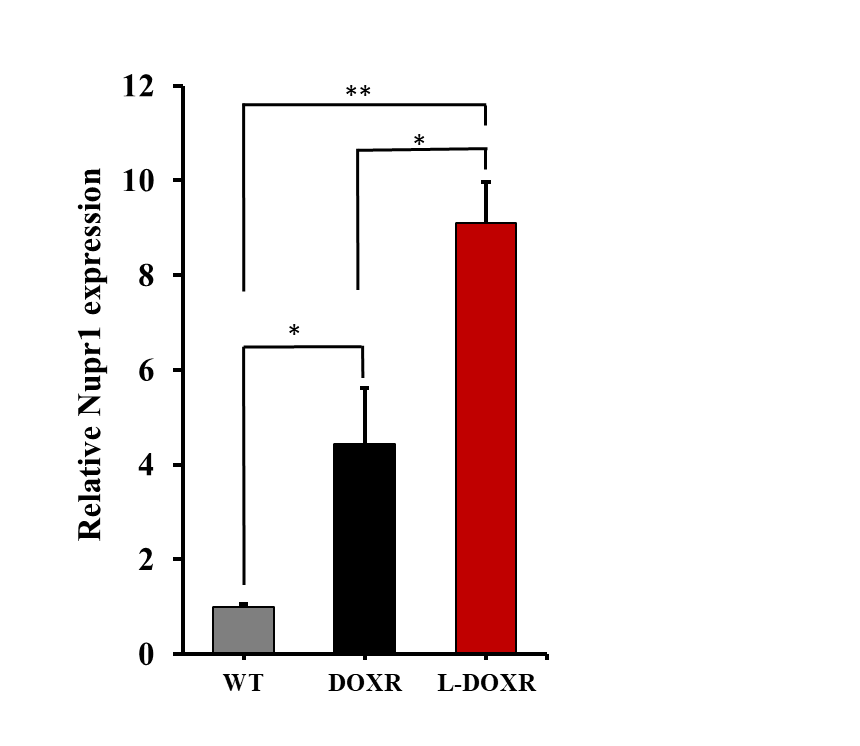

**Supplementary Fig. 3 (related to Fig. 4). *Nupr1* expression in WT, DOXR and L-DOXR.** The RNA expressions of *Nupr1* were measured by RT-qPCR. All data are presented as means ± SEM; ***p < 0.001; Student’s two-tailed, unpaired t-test.

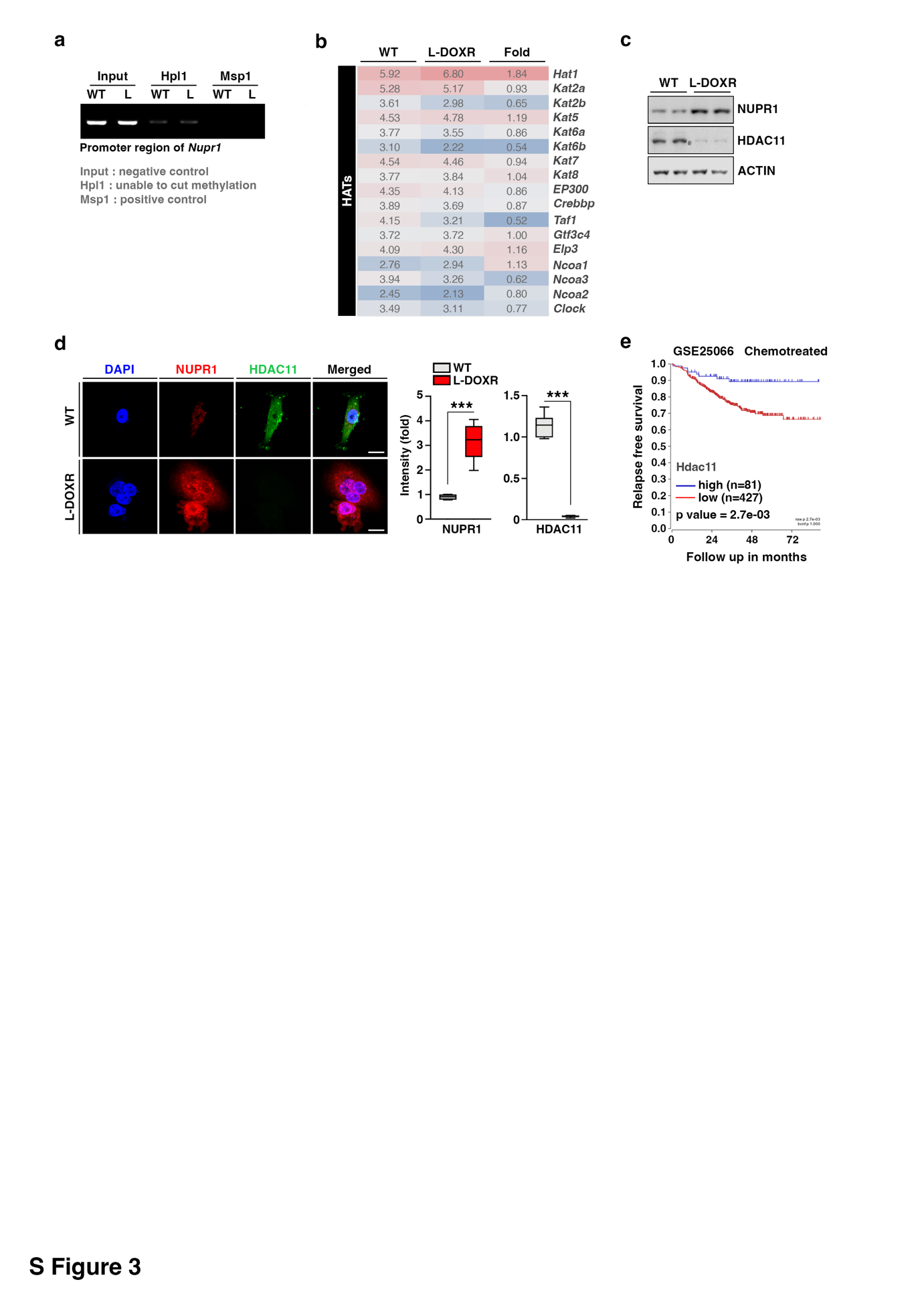

**Supplementary Fig. 4 (related to Fig. 5). HDAC11 suppression leads to *Nupr1* upregulation in L-DOXR.** (a) Extracted gDNA from WT and L-DOXR were digested with restriction enzymes that cut only non-methylated CCGG or cut all CCGG sequences. gDNA was assessed for methylation levels using RT-PCR with specific primer sets for CpG islands of *Nupr1*. L, PACCs. (b) Heatmap representing the relative mRNA expression levels of *Hdacs* in WT or L-DOXR. (c) Protein expressions of NUPR1 and HDAC11 were measured from WT and L-DOXR by immunoblotting. (d) Cells were stained with anti-NUPR1 (red) and HDAC11 (green) antibodies for measuring their negative correlation. The intensity of NUPR1 and HDAC11 was measured by ImageJ. Scale bar: 20 μm. (e) Comparison of overall survival between high (n=81) or low (n=427) expressions of Hdac11 in patients after chemo treated (GSE25066, n=508) were analyzed by the Kaplan-Meier and Log-rank test. All data are presented as means ± SEM; ***p < 0.001; Student’s two-tailed, unpaired t-test (d).

**table S1. List of primer sequences.**

| **ChIP assay primers** | | |
| --- | --- | --- |
| **Gene Name** | **Forward Primer** | **Reverse Primer** |
| Nupr1 -1400/-1000 | GATCTCAGCTCACCACAACCT | ATCTTCTTCCTAGAGTTGGGGAGA |
| Nupr1 -1000/-600 | ACAATTATTATCATCCTTATTTTACAG | ATAGACATCTGCCACCATGCC |
| Nupr1 -600/-200 | AATCCCAGCTATTCGGGAGG | ATATTTTCCATAGAGGAGGTCCCG |
| Nupr1 -200/+200 | CCAGCTGGGTGAGCCTGG | AGATGGCTGAGTGGGCCTTA |
| **Real-time qPCR primers** | | |
| **Gene Name** | **Forward Primer** | **Reverse Primer** |
| Gapdh | GTGTTCCTACCCCCAATGTGT | ATTGTCATACCAGGAAATGAGCTT |
| Nupr1 | TCGGGCCTCTCATCATGCCT | TGCCCCTCGCTTCTTCCTCT |
| Hdac1 | ATATCGGGGCTGGCAAAGGC | TCCACACACTTGGCGTGTCC |
| Hdac2 | AGGCCCCATAAAGCCACTGC | CTCCAGCAACTGAACCGCCA |
| Hdac3 | CTCAGCATCCGAGGGCATGG | TCGATGCGGGTGCTGACATC |
| Hdac4 | GGTGGTGTTGGGGTGGACAG | GCTCTCCTCCGCATGGTGTC |
| Hdac5 | CAACAGCTCCCACAGCACCA | TTGGTGACAGTGACCGTGGC |
| Hdac6 | GACCTTGGAGCTAGGCAGCG | TGCCACCAAATGGGGACACC |
| Hdac7 | CAGTGACCGCAGGACCCATC | CTGGGCAAAGTGGAAGGGCA |
| Hdac8 | CAACACGGCTCGATGCTGGA | CGGCAGCTTGGCGTGATTTC |
| Hdac9 | TATGGCACCAACCCCCTGGA | GCACCGGACGAGTGTAGCTC |
| Hdac10 | CCTGGCCTATGGCTTCCAGC | TGCTAGCTGGGGTGTGGAGT |
| Hdac11 | GTGCACACGAGGCGCTATCT | GCCAGCTTCCCCGCCATTAT |
